# Deficits in seizure threshold and other behaviors in adult mice without gross neuroanatomic injury after late gestation transient prenatal hypoxia

**DOI:** 10.1101/2021.08.04.451528

**Authors:** Ana G. Cristancho, Elyse C. Gadra, Ima M. Samba, Chenying Zhao, Minhui Ouyang, Sergey Magnitsky, Hao Huang, Angela N. Viaene, Stewart A. Anderson, Eric D. Marsh

**Author notes:** Corresponding Author: Ana G. Cristancho, MD, PhD, Abramson Research Center, Rm. 516g, Children’s Hospital of Philadelphia, 3615 Civic Center Blvd., Philadelphia, PA. 19104, U.S.A.

## Abstract

Intrauterine hypoxia is a common cause of brain injury in children resulting in a broad spectrum of long-term neurodevelopmental sequela, including life-long disabilities that can occur even in the absence of severe neuroanatomic damage. Postnatal hypoxia-ischemia rodent models are commonly used to understand the effects of ischemia and transient hypoxia on the developing brain. Postnatal models, however, have some limitations. First, they do not test the impact of placental pathologies on outcomes from hypoxia. Second, they primarily recapitulate severe injury because they provoke substantial cell death, which is not seen in children with mild hypoxic injury. Lastly, they do not model preterm hypoxic injury. Prenatal models of hypoxia in mice may allow us to address some of these limitations to expand our understanding of developmental brain injury. The published rodent models of prenatal hypoxia employ multiple days of hypoxic exposure or complicated surgical procedures, making these models challenging to perform consistently in mice. Furthermore, large animal models suggest that transient prenatal hypoxia without ischemia is sufficient to lead to significant functional impairment to the developing brain. However, these large animal studies are resource-intensive and not readily amenable to mechanistic molecular studies. Therefore, here we characterized the effect of late gestation (embryonic day 17.5) transient prenatal hypoxia (5% inspired oxygen) on long-term anatomical and neurodevelopmental outcomes in mice. Late gestation transient prenatal hypoxia increased hypoxia-inducible factor 1 alpha protein levels (a marker of hypoxic exposure) in the fetal brain. Hypoxia exposure predisposed animals to decreased weight at postnatal day 2, which normalized by day 8. However, hypoxia did not affect gestational age at birth, litter size at birth, or pup survival. No differences in fetal brain cell death or long-term gray or white matter changes resulted from hypoxia. Animals exposed to prenatal hypoxia did have several long-term functional consequences, including sex-dichotomous changes. Hypoxia exposure was associated with a decreased seizure threshold and abnormalities in hindlimb strength and repetitive behaviors in males and females. Males exposed to hypoxia had increased anxiety-related deficits, whereas females had deficits in social interaction. Neither sex developed any motor or visual learning deficits. This study demonstrates that late gestation transient prenatal hypoxia in mice is a simple, clinically relevant paradigm for studying putative environmental and genetic modulators of the long-term effects of hypoxia on the developing brain.

## Introduction

Intrauterine hypoxia is one of the most common causes of long-term brain injury worldwide [1]. The most immediately clinically evident form of prenatal hypoxia is neonatal hypoxic-ischemic encephalopathy (HIE) in term infants due to an intrapartum loss of oxygen and nutrients. HIE impacts millions of births each year, with about 40% of affected children having a neurodevelopmental disability (NDD) such as autism and epilepsy [1, 2]. Children initially classified as having mild HIE, defined as minimal evidence of perinatal cell death by imaging and relatively normal neonatal physical exam, may still experience adverse outcomes [3–8]. These children do not qualify for therapeutic hypothermia, the only specific intervention for moderate to severe HIE [9]. In addition to term-infant HIE, preterm HIE is a poorly understood entity that also predisposes children to NDDs [10, 11]. Most infants with preterm HIE are also not eligible for therapeutic hypothermia. Since the incidence of HIE is disproportionately higher in children born in areas with limited prenatal care or access to therapies [12], there is an urgent need for novel, widely available interventions.

Most rodent studies investigating the effects of transient hypoxia on the developing brain occur in the postnatal brain (day of life 7 to 10) [13–15]. Postnatal models have provided many insights, in part because the rodent brain is neuroanatomically considered human “term” equivalent at postnatal days 7 to 10 [16] (**Fig. 1A**). Postnatal studies are also more technically amenable to studying the combined effects of hypoxia and ischemia [13–15]. However, it is likely that to understand the impact of prenatal hypoxia on the developing brain, we must consider that *in utero* hypoxic injury likely <u>differs</u> from postnatal injury. The *in utero* environment is relatively hypoxic at baseline [17]. The required oxygen tension during fetal development is tightly regulated, as evidenced by brain injury from both early exposure to relative hyperoxia in premature children or *in utero* hypoxia [18, 19].

**Fig. 1:**
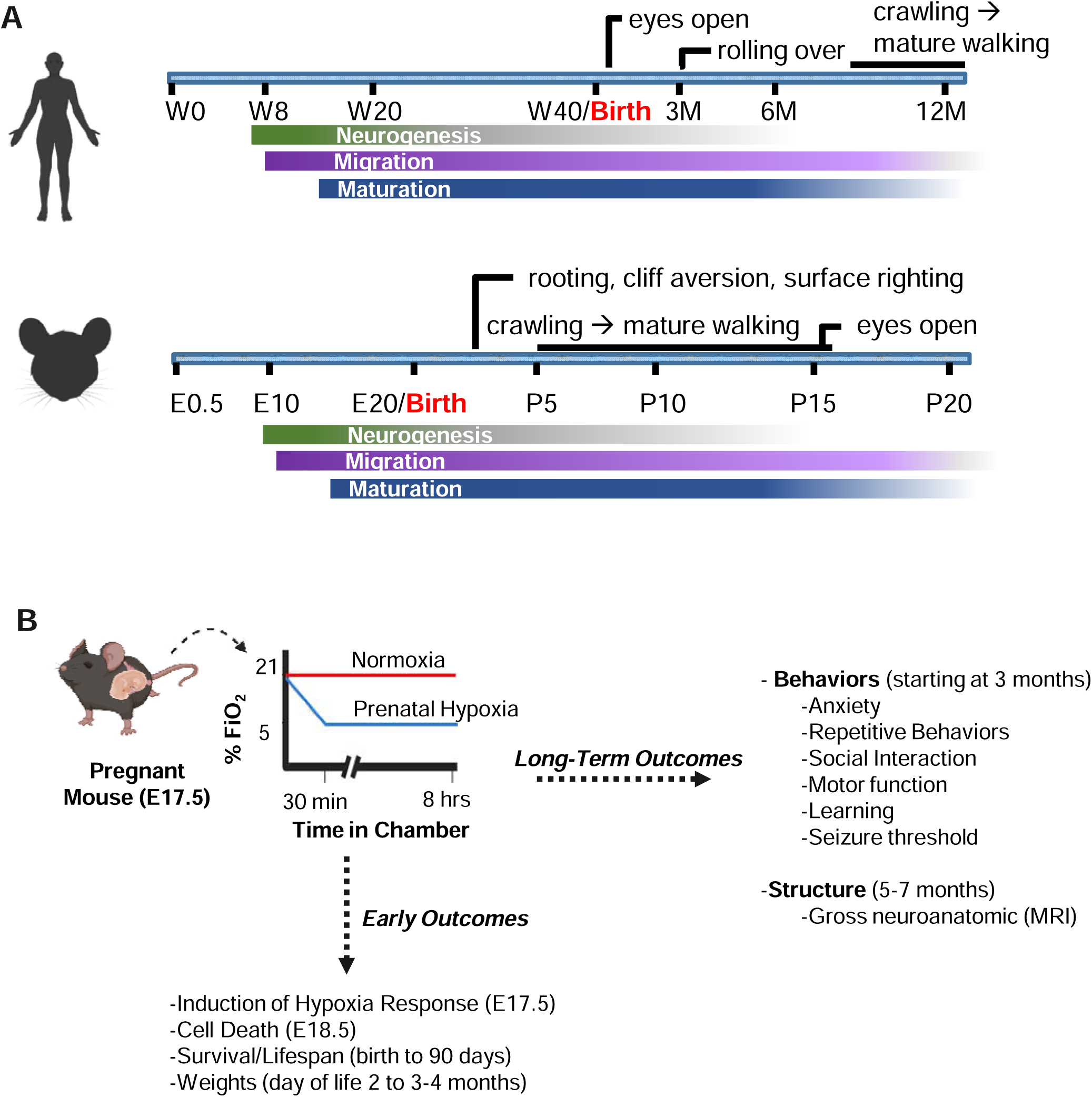
Comparison of human and mouse neuroanatomic and milestone development with a schematic of prenatal hypoxia model. (A) Timeline of human and mouse brain development. The human timeline spans the weeks of gestation (W) and months after delivery (M). The mouse timeline includes embryonic age (E) and postnatal days of life (P). Specific developmental milestones are indicated. (B) Schematic and timing of late gestation transient prenatal hypoxia exposure and performed experiments.

In response to the intrinsically hypoxic environment *in utero*, the protein hypoxia-inducible factor 1 alpha (Hif1α) is stabilized in the fetal brain [20]. Hif1α is a transcription factor required for *in utero* neuronal development [21] and plays a vital role in oligodendrocyte differentiation [22]. Hif1α induces the expression of many target genes to orchestrate the initial adaptive response to hypoxia [20]. There are conflicting studies on the role of Hif1α in HIE [23–25]; it has been described as both neuroprotective and neurotoxic. This discrepancy in the role of Hif1α may be partly explained because it has generally been studied in the postnatal environment and not when it is stabilized in the intrauterine environment. The literature suggests differences in the intrinsic oxygen levels in the prenatal and postnatal brain may lead to contrasting responses to hypoxia. Hypoxic preconditioning in postnatal rodent models of HIE is highly protective of severe injury [25, 26]. Conversely, prenatal hypoxia decreases the uterine and placental blood flow within minutes [27], and maternal conditions partially associated with placental hypoxemia worsen clinical outcomes from HIE [28].

There are other limitations to postnatal models for mild HIE and preterm HIE. For example, in the most common model using a carotid ligation followed by hypoxia at postnatal days 7 to 10 (known as the Vannucci model) [13], there is significant cell death, most consistent with severe damage, and focal injury, most analogous to perinatal stroke [13, 29]. HIE, however, is a global insult to the entire brain and can still result in significant neurodevelopmental impairment even without evidence of significant cell death on perinatal imaging [3–8]. Additionally, functional networks do not correlate with neuroanatomic maturation in rodents when compared with human development [30] (**Fig. 1A**). At birth, mice are considered neuroanatomically equivalent to infants born at 23-26 weeks gestation [16]. Yet, they do not have the respiratory or feeding dyscoordination at birth that is characteristic of children born before 34 weeks gestation [31]. In addition, mice have an immature walk by postnatal day 10, a developmental milestone that infants reach at about 12 months of age [30, 31]. Lastly, by definition, postnatal models are not designed to model the effects of hypoxia on the premature brain. Therefore, using transient prenatal hypoxia models may help us understand key aspects of the pathophysiology of HIE as they affect *in utero* brain development.

Mouse models of prenatal hypoxia exist, but they primarily study the effects of chronic hypoxia by exposing dams to mild hypoxia (~ 10% inspired oxygen) throughout gestation [32, 33] or are technically challenging (late gestation uterine artery ligation) [34]. Nevertheless, some studies of the long-term effects of transient hypoxia in more simplified models are published. For instance, adult mice demonstrate increased anxiety-like behaviors when exposed prenatally to a hybrid hypoxia model with elevated carbon dioxide (2 hours of 9% inspired oxygen with 3% inspired carbon dioxide) [35, 36]. Otherwise, most of our understanding of brief, mild hypoxia-only insults comes from large animal studies. For example, in sheep, when compared with a model of hypoxia plus ischemia, transient prenatal hypoxia alone leads to long-term structural and functional injury despite less cell death than the ischemic condition [37, 38]. The deficits from hypoxia alone included decreased dendritic complexity and abnormal action potential propagation [37, 38]. Unfortunately, these experiments in sheep require specialized facilities, extensive resources, and are not amenable to genetic manipulation for mechanistic studies. Finally, existing studies of transient prenatal hypoxia have not addressed the potential of sex-specific differences in response to prenatal brain injury, which is described in postnatal models [39–42].

Therefore, here we characterize a simplified model of late gestation transient prenatal hypoxia in male and female mice. Our goal is to determine whether this model is amenable to future mechanistic studies of the maternal-placental-fetal unit and if it may serve as a preclinical model of mild HIE and preterm HIE. In particular, we focus on early and late neuroanatomic injury using clinically anaolgous pathology and neuroimaging measures and describe a comprehensive battery of behavioral studies to determine whether this model correlates to mild hypoxic injury seen in children.

## Materials and Methods

### Animals

Male and female C57Bl/6 mice were purchased from Charles River Laboratories (Wilmington, MA). Timed matings were employed to ensure consistency for gestational age of prenatal hypoxia exposures. The mice were maintained at the Children’s Hospital of Philadelphia Animal Facility at a 12-hour light, 12-hour dark cycle (0615-1815 h), and with *ad libitum* access to water and diet mouse food 5015 (LabDiet, St. Louis, MO). Offspring exposed to prenatal normoxia or hypoxia were weighed at postnatal day 2 (P2), P8, and before all behavioral assays by experimenters blinded to exposure conditions. Animals were group-housed for postnatal experiments. Male and female gonadectomized CD1 mice for Social Interaction were purchased from Charles River Laboratories. The Institutional Animal Care and Use Committee of the Children’s Hospital of Philadelphia approved all experiments.

### Prenatal hypoxia

Pregnant females were placed in a controlled oxygen chamber (BioSpherix Ltd., Parish, NY) at embryonic day 17.5 (E17.5, **Fig. 1B**). Mice were acclimated to the chamber for 1-2 minutes at 21% inspired oxygen. For hypoxic exposures, oxygen concentration was decreased with balanced nitrogen to 5% inspired oxygen over 30 minutes and maintained at 5% inspired oxygen for the duration of the experiment (2, 4, 6, or 8 hours). Pregnant mice were placed in the same controlled oxygen chamber for 8 hours for the normoxic control exposures at 21% inspired oxygen. Dams in both groups were provided free access to food and hydrogel for hydration throughout the time in the chamber. For locomotion, dams (normoxia n=4, hypoxia n=4) were videotaped during the first 90 minutes in the chamber. We quantified the cumulative distance the mouse moved in the chamber using ANY-maze software (Stoelting Co., Wood Dale, IL, U.S.A.). Except for protein and RNA studies, pregnant dams were monitored for recovery after hypoxia and demonstrated normal locomotion within 10 minutes of the end of the exposure. All survival studies were performed after 8 hours of prenatal hypoxia (normoxia n = 19, hypoxia n = 28.) Pregnant dams from normoxia and hypoxia were assessed twice per day until delivery to determine gestational age at birth and the number of pups per litter at birth.

### Protein isolation and immunoblot

Immediately after 2, 4, 6, or 8 hours of hypoxia or 8 hours of normoxia, pregnant mice were anesthetized with isoflurane, euthanized with a cardiac puncture, and whole fetal brains were harvested from the offspring. Brains were hemi-dissected; one-half was used for protein isolation and the other half was used for RNA isolation. Samples were immediately flash-frozen in liquid nitrogen for future processing. For protein isolation, a Biopulverizer was cooled with liquid nitrogen and used to pulverize the tissue before the addition of RIPA buffer with phosphatase inhibitors (cOmplete, Mini, Sigma, St. Louis, MO) and sonication (n=4 mice/condition, each mouse in a condition was from a separate litter). The resulting solution was centrifuged at 4°C for 15 minutes at 14,000 g and the supernatant was collected. Total protein in the lysate was quantified via a BCA assay (Thermo Scientific, Rockford, IL). A 4 to 20% gradient Tris-Glycine gel was loaded with 30 μg of protein. The gel was transferred to a nitrocellulose membrane using the Invitrogen iBlot system (Thermo Fisher, Waltham, MA) and blocked in 5% milk/1x TBS for 60 minutes at room temperature. Blots were incubated overnight at 4 °C in polyclonal rabbit anti-Hif1α antibody (Novus Biologicals, Littleton, CO; NB-1000-134, 1:1,000). Blots were washed 3 times in 5 minutes in TBST (Tris-buffered saline + 0.1% Tween-20) and incubated in fluorescent secondary antibody for 1 hour at room temperature before imaging (donkey anti-rabbit; Licor, Lincoln, NE; 1:5,000). Blot was probed for actin using an anti-β-actin antibody (1:5,000) and fluorescent secondary. Blots were quantified using ImageJ.

### RNA isolation and quantitative PCR

We isolated total RNA from flash-frozen whole brain samples using the RNeasy Lipid Tissue Mini Kit (QIAGEN, Venlo, Netherlands). RNA was isolated with DNAse to avoid genomic DNA contamination. Individual samples shown were isolated from n =11-12 fetal brains/condition in 3 separate litters/condition. We converted 2 μg of RNA to cDNA with the High-Capacity cDNA Reverse Transcription Kit (Applied Biosystems, Foster City, CA). Quantitative PCR on Quant Studio 12K Flex (Applied Biosystems, Foster City, CA) was used to calculate mRNA levels. *Vegfa* mRNA levels (Taqman, Mm00437306_m1, Thermo Fisher Scientific, Waltham, MA levels, a HIF1α target, were normalized to 18S mRNA levels (Taqman, Mm03928990_g1, Thermo Fisher Scientific, Waltham, MA) in each sample. To normalize between different plates, we standardized the average of the normoxic condition to 1, and all other samples on that plate were multiplied by the sample standardization factor.

### Maternal nestlet

The ability for pregnant dams to form a nest after normoxia or hypoxia was used as a proxy for the ability to create a nurturing environment. A 5-point scale was utilized as previously described, with a score of 5 being an intact nest [43]. In brief, after 8 hours in the oxygen chamber in normoxia or hypoxia, pregnant dams were moved to a new cage with an intact cotton square and no other cage enrichment. They were placed in a standard holding room overnight. The following morning an experimenter obtained an overhead picture of the cage with minimal disruption. A blinded experimenter scored the images.

### Histology

Fetal brains were harvested 24 hours after exposure and placed in 4% paraformaldehyde. Fixed brains were processed and paraffin-embedded and then sectioned in the coronal plane by the CHOP Pathology Core into 5 μM unstained slides. Every fifth slide was stained with hematoxylin and eosin and examined to identify brain regions of interest. Then, the apoptotic nuclei in the cortex, basal ganglia, and white matter were counted by a neuropathologist (A.N.V.), who was blinded to the sample condition.

### Ex vivo MRI

High-resolution diffusion tensor imaging (DTI) is an effective noninvasive imaging tool to delineate neuroanatomy [44–46]. We used a high-resolution DTI (0.1×0.1×0.1mm) with an MR scanner of high magnetic strength to examine the morphological and microstructural changes in mice with hypoxia. MRIs for 7 normoxic animals (5 females, 2 males) and 6 hypoxic animals (4 females, 2 males) were analyzed. Animals were 5-7 months old at the time of the MRIs (2-3 weeks after completion of the flurothyl studies).

Mice were anesthetized with 5% isoflurane. We then performed trans-cardiac perfusion with chilled phosphate-buffered saline (PBS, Sigma St. Louis, MO) for 5 min at 8 mL/min then with 10 % buffered formalin (Sigma, St. Louis, MO) and 10 % of 0.5 mmol/L of gadoteridol (Gadavist, Bayer Healthcare Pharmaceuticals, Whippany, NJ or Prohance, Bracco Diagnostics, Princeton, NJ) in PBS for another 5 min at 8mL/min. Next, we removed the skin from the skull, and the heads were submerged in 10% buffered formalin overnight. The following day we detached the muscle, eye, and mandible from the skull, and the head was placed in 0.5% of 0.5 mmol/L of gadoteridol in PBS for 7 days. Skulls were then placed in PBS for at least 24 hours before the MRI. Then, we transferred the skulls into an MR-compatible tube and immersed them in fomblin (Fomblin Perfluoropolyether; Solvay Solexis, Thorofare, NJ) during the MRI scan.

### MRI data acquisition

The MRI data were acquired on a Bruker 9.4T vertical bore scanner. A volume coil with an inner diameter of 15mm was used as an RF transmitter and receiver. A 3D high-resolution, multi-shot echo-planar imaging (EPI) sequence with eight shots was used to acquire diffusion-weighted images (DWIs). The parameters for DWIs were as follows: field of view=25.6×12.8×10.0mm; voxel size=0.1×0.1×0.1mm; echo time=26ms; repetition time=1250ms; 6 or 30 independent diffusion-weighted directions with a b value of 1500 s/mm^2^ and five additional images without diffusion gradients; two averages.

### Diffusion tensor fitting

Diffusion tensor was fitted in DTIStudio (http://www.MRIstudio.org) [47]. After the diagonalization of the tensor to obtain three eigenvalues (λ_(1-3)) and eigenvectors (ν_(1-3)), mean diffusivity (MD) was calculated as the mean of three eigenvalues (λ_(1-3)). Fractional anisotropy (FA) was calculated as follows.

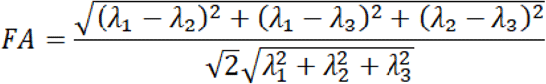

### Regions of interest (ROIs) delineation and measurements of volume and thickness

The experimenter was blinded to the exposure condition for each animal when defining all the ROIs. The ROIs, including lateral ventricles, were manually placed on the averaged b0 map or MD map by referencing known stereotactic reference atlas [48] in ROIEditor (http://www.MRIstudio.org). For genu and splenium of the corpus callosum, the ROIs of consistent size were drawn on seven slices centered around the mid-sagittal slice of the brain (the mid-saggital slice plus 3 slices on the left and right of that central slice). The volume of the ventricles was calculated as the number of voxels in the ROI times the voxel size. In addition, we measured the thickness of the anterior cingulate cortex in the mid-sagittal region. Volume and thickness measurements were calculated in Matlab (Mathworks, Inc., Natick, MA, U.S.A.).

### Behavior

Behavior studies following published protocols were performed sequentially starting at 3 months old in the order listed below for all animals [49–55]. Any modifications are listed below. Three separate cohorts consisting of 5 normoxic litters and 4 hypoxic litters were used. Composition of experimental groups consisted of normoxic male = 16, hypoxic male = 10, normoxic female = 22, hypoxic female = 13.

All of the testing except for grip strength was performed between 0700 h to 1300 h. Grip strength was conducted from 1400 h to 1600 h. All animals were given 1 to 7 days of rest between each behavior paradigm to minimize any carrying effects between the tests. Animal weight was monitored before each test as a proxy of health (no mice were removed from the testing due to weight loss over 10% or any other visible markers of distress.) Animals were not acclimated to handling before elevated zero maze or open field but were subsequently handled daily between experimental days for 5 minutes per cage. Mice were adapted to the testing room for at least 30 minutes before experimentation. We cleaned the equipment apparatuses between mice (PDI Sani-Cloth Plus germicidal disposable cloth). ANY-maze software (Stoelting Co., Wood Dale, IL, U.S.A.) was used to video-record and score elevated zero maze, open field, social interaction, and Morris water maze.

### Elevated zero maze

The elevated zero maze test was modified from a previous description [55]. In brief, an annular 60 cm diameter beige apparatus was elevated 50 cm above the ground, with care to ensure it was level. Two opposing quadrants were “open,” where the mouse could freely peer over the edge of the ring. The other two opposing quadrants were “closed,” with 16 cm opaque walls preventing the animal from seeing over the edge. The experimenter was hidden from the animals in the apparatus by a black drape. Illumination of the apparatus was standardized for all the cohorts to 150-200 lumens/m^2^ in the open arms and 50 lumens/m^2^ in the closed arms. Animals were placed in the center of a closed arm and allowed to roam the apparatus freely for 5 minutes. ANY-Maze was used to calculate the animals’ time in the open and closed arms.

### Open field

The open field test was modified from a described protocol [54]. In brief, A 53 × 53 cm white plastic box with 22-cm high walls and no top was placed on the floor. Illumination was from overhead fluorescent lights in the 150-200 lumens/m^2^ range. The experimenter was visually separated from the animals in the apparatus by a black drape. Each mouse was placed in the center of the box. The mouse was then allowed to roam freely in the box for 15 min. ANY-maze was used to calculate the animal’s time in the periphery or center of the box. The periphery was defined as within 13 cm of the box wall. Otherwise, the animal was described as being in the box’s center. The distance traveled during the test in the entire apparatus was also measured.

### Marble burying

The marble burying test was modified from previous studies [49, 51]. A white 41.9 cm × 33.7 cm × 22.5 cm box was filled with 5 cm of fresh bedding. Twenty-four marbles were arranged in a 4 × 6 grid 5 cm apart. Up to 6 mice were tested simultaneously, so barriers were placed between the boxes. Overhead lights provided illumination with a range of 130-150 lumens/m^2^ at each box. Each mouse was placed in the center of the box for 30 minutes and allowed to roam freely. The experimenter was visually separated from the animals in the apparatus by a black drape. Pictures were taken of each box before the animal was placed inside and after the mouse was carefully removed at the end of the experiment. Two blinded reviewers examined images to determine whether marbles were buried greater than two-thirds into the bedding. Average from the two reviewers of marbles buried for each mouse is reported.

### Short-term nestlet

A short-term nestlet experiment was modified from a previous study [51]. Animals were placed in a clean cage with 1 cm of fresh bedding. A precut nestlet with all rough edges removed was weighed and placed in the center of the cage. Each mouse was placed in the chamber’s center, facing away from the experimenter. Then, the cage lid was placed on top. The mouse was allowed to roam freely in the chamber for 30 minutes. As multiple animals were in the cages simultaneously, we put barriers between the cages to visually separate the animals. The experimenter was positioned out of sight of the animals. At the end of the 30 minutes, the animals were removed. Nestlets were removed from the cage and allowed to dry overnight. A blinded experimenter handled all the neslets and weighed them after > 24 hours of drying. The change in nestlet weight was calculated by subtracting from the pre-experiment nestlet weight.

### Rotarod

Mice were tested on a Rotarod from Ugo Basile (Model 47650, Comerio, Italy) as previously published [53]. The mice were tested in groups of up to 5 animals for each trial. For 4 days, we performed 3 trials per day, each a maximum of 5 minutes long. The Rotarod was used in acceleration mode. In the first 2 days, the speed increased from 4 rpm to 40 rpm. In the last 2 days, the speed increased from 8 rpm to 80 rpm. We recorded the rate (speed) when the mouse fell off the rod or if the mouse was grappling the rod without walking for 4 consecutive cycles. If the mouse was still walking well on the rod at the end of 5 minutes, then the maximum speed of the trial was recorded and we stopped the trial. The mice were placed back into individual containers between the trials and then put back in home cages at the end of the last trial. The intertrial time was 15-20 minutes.

### Social interaction

Social interaction testing was performed as previously described [54]. A white three-chamber apparatus was filled with 1 to 2 cm of fresh bedding. This experiment was performed in red light without fluorescence to resemble a night environment where mice are most likely to interact. The experimenter was visually separated from the animals in the apparatus by a black drape. For the habituation stage, empty clear tubes with holes were placed in the center of the left and right chambers. The experimental mouse was then allowed to freely roam the apparatus for 10 minutes. Then, the experimental mouse was placed in a temporary holding box to reset the apparatus for the next stage. For the novel mouse/object stage, a sex-matched gonadectomized mouse was placed in the tube on the left. A novel object (a small featureless toy) was placed in the tube on the right. The experimental animal was then returned to the apparatus to roam freely for 5 min. The time in each chamber (left, right, and center) was recorded in ANY-Maze for both stages.

### Grip strength

The grip strength of the mice was tested using a grip strength meter (Model 080312-3 Columbus Instruments, Columbus, OH, U.S.A.) as previously published by the Marsh lab [54]. In brief, each mouse was tested in 6 consecutive trials; 3 for forelimbs and 3 for hindlimbs. In each trial, limb strength was recorded as kilograms of force. The average strength for each mouse for forelimb and hindlimb trials is reported.

### Morris water maze

As previously published, the Morris water maze test was performed in a 128 cm round plastic tube filled with room temperature water (approximately 21°C) [50, 54]. The testing was performed in 3 phases: visual acuity trials, place trials, and probe testing. For the first two phases, a platform was submerged 0.5 cm below the water surface and the water was opacified with non-toxic white tempera paint so that the platform was not visible to the animal. A mouse could swim for up to 60 sec to find the platform and was led to the platform if it could not locate it by 60 sec. Once on the platform, the mouse was maintained on the platform for 15 seconds. The mouse was then removed from the platform, dried, and placed in an individual container under a heat lamp in between trials. There were 4 trials per day.

Specifically, in the first phase, the visual acuity of each mouse was tested for 2 days by attaching a flag that was visible above the platform. Then, the platform and flag were placed in a different quadrant and the mouse was started in a different position for each trial. A mouse was considered to have adequate vision if by the 2nd day it was able to find the platform at least 50% of the trials before 60 seconds. All mice were deemed to have adequate vision to continue with the next phase.

The place trials were performed over 5 days. We placed visual cues in regions labeled as “north,” “south,” “east, “and “west” in the tub. The flag was removed from the platform, and the platform was completely submerged in the southwest quadrant. For each trial, a mouse was placed in a different location of the tub. The time to platform was measured in ANY-Maze, and the average of these measures for all the trials for each day was reported.

For probe testing, the platform was removed from the pool. One hour after the last place trial, each mouse was placed in the northeast quadrant and allowed to swim freely for 60 seconds to assess short-term memory. The time spent swimming in the southwest quadrant, where the platform was previously located, was measured. Next, twenty-four hours after the last place trial, the probe test was repeated to assess long-term memory.

### Flurothyl seizure threshold

Flurothyl seizure threshold testing was modified from a previously described protocol [56, 57]. Flurothyl (bis-2,2,2-trifluoroethyl ether, Sigma, St. Louis, MO) was diluted in 100% ethanol to make a 10% flurothyl/EtOH solution. In brief, 5 to 6-month-old mice were placed on a platform in a 1.7 L rubber-sealed glass chamber (Ikea, Älmhult, Sweden) containing a small amount of the carbon dioxide scavenger soda-lime (Sigma). The flurothyl solution was infused with a syringe pump at a 6 mL/hour rate until the first generalized tonic-clonic seizure (GTC) was noted. When the first GTC was noted, the pump was stopped and the chamber lid was removed. The animals were videotaped throughout the experiment. The experimenter was blinded to the exposure condition during the experiments. Two blinded scorers used a modified Racine scale to define a generalized tonic-clonic seizure [15].

### Statistical analysis

All animal experiments were performed with the experimenter blinded to the exposure condition. Most of the data is presented using truncated violin plots with individual mice plotted superimposed on top to display frequency distribution (GraphPad, San Diego, CA). The data are grouped by condition and sex (when sufficiently powered). Markings on violin plots are as follows: the center dashed line is the median value, and the lighter lines demonstrate quartiles. We did not remove any mice from the analysis as outliers even if they met the statistical criteria for outliers. Since we did not identify any experimental anomalies that could account for the differences, we reasoned that the variability in any experiment is more likely a reflection of the biological variability due to hypoxia exposure.

For statistics, we used GraphPad and R-studio. The area under the curve was calculated to quantify the locomotion of pregnant dams, and the unpaired t-test was used to test for statistical significance. The Mann-Whitney test was used to compare normoxia and hypoxia maternal nestlet shredding, gestational age, litter size, and MRI studies. The log-rank test was utilized for survival. For Hif1α protein levels, the One-Way ANOVA was used. For *Vegfa* mRNA, the One-Way Nested ANOVA was used in Graphpad. Dunnett’s multiple comparison test was used after both One-Way ANOVA tests. Welch’s t-test was used to compare apoptosis between normoxia and hypoxia.

For all other studies, R was used for statistics. We utilized mixed models with generalized estimating equations in the geepack package in R for most of the comparisons [58]. Linear mixed-effects models in the lme4 package in R was used for experiments requiring analysis of daily repeated measures (weights, Rotarod, Morris Water Maze) [59]. The cohort was used as a random effect. Independent variables for all models included prenatal hypoxia exposure, sex, gestational age at birth, and litter size at birth. Statistical significance was displayed for hypoxia, sex, and interaction between hypoxia and sex since hypoxia has been shown to have sex-dichotomous effects [60, 42]. Statistical significance was set at * *p* < 0.05, ** *p* < 0.01, and *** *p* < 0.001. A Benjamini-Hochberg correction was used for all *p*-values obtained from mixed model analyses. Statistical subanalysis of sex-specific differences for hypoxia was only performed if hypoxia or interaction between sex and hypoxia had an adjusted *p* < 0.15. These criteria allowed us to determine if there were more subtle effects unique to each sex due to hypoxia exposure that may not have been evident when all the animals were analyzed together. The Shapiro-Wilk test was used to determine the normality of the distribution within each test. If the distribution was normal, the mean and standard deviation were reported. The median and interquartile range (IQR) was reported if the distribution was not normal.

## Results

### Prenatal hypoxia induces the canonical hypoxic response in the fetal brain

First, we performed a time course experiment to determine the duration of prenatal hypoxia required to induce a reproducible hypoxic molecular response in the fetal brain. The transcription factor Hif1α and its canonical target, vascular endothelial growth factor A (*Vegfa*), are established molecular markers of the hypoxic response [61]. Hif1α protein is stabilized by the intrinsically hypoxic intrauterine environment [20], and prenatal hypoxia can further stabilize this protein **(Fig. 2A & 2B)** (mean/SD normoxia = 0.11 +/−0.09, 2 hours = 0.28 +/−0.18, 4 hours = 0.42 +/−0.15, 6 hours = 0.29 +/−0.11, 8 hours = 0.24, +/−0.13). *Vegfa* mRNA levels also increased with prenatal hypoxia **(Fig. 2C)** (mean/SD normoxia = 1 +/−0.30, 2 hours = 2.26 +/−0.73, 4 hours = 2.62 +/−0.81, 6 hours = 2.93 +/−1.24, 8 hours = 2.50, +/−0.48). While the peak increase for both protein and mRNA was at 4 hours of hypoxia, there was decreased variability in *Vegfa* after 8 hours of hypoxia. Thus, we chose to expose the dams to 8 hours of hypoxia for all further experiments.

**Fig. 2:**
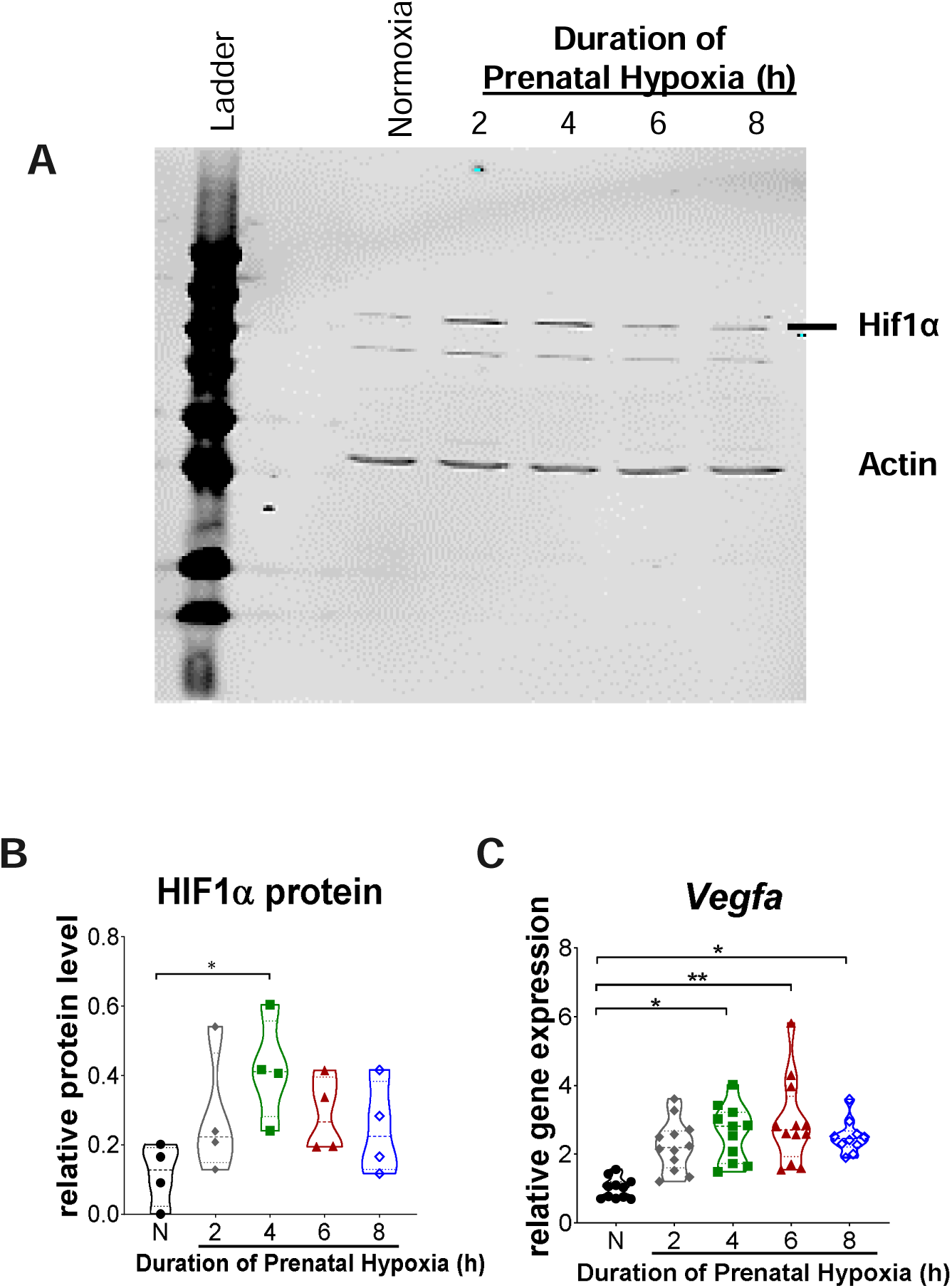
Prenatal hypoxia paradigm induces a canonical HIF1α response consistent with hypoxic insult in the fetal brains. (A) Representative immunoblot if HIF1α (~120 kDa) and actin (~40 kDa) in fetal brains after the indicated duration of prenatal hypoxia. (B) Quantification of HIF1α protein level as normalized to actin, the loading control. (C) Vegfa mRNA levels in the fetal brain in normoxia and after the indicated duration of hypoxia. The points shown in the graphs are from individual fetal brains. Statistics are described in the methods section.

### Maternal and offspring survival and health were unaffected by prenatal hypoxia

Pregnant mice demonstrated decreased locomotor activity during hypoxic exposure once the oxygen level was less than 10% inspired oxygen, approximately 20 minutes after beginning the hypoxia exposure (the endpoint of 5% inspired oxygen was reached at 30 minutes) **(Fig. 3A)** (mean/SD area under the curve first 20 min: normoxia = 19.5 +/−5.2, hypoxia = 18.1 +/−4.0; area under the curve last hour: normoxia = 221.0 +/−59.8, hypoxia = 107.2 +/−16.4). The pregnant mice survived 8 hours of hypoxia and rapidly recovered within 10 minutes of the end of the hypoxia exposure. Since we saw a decrease in maternal locomotion with hypoxia, we tested whether the maternal dam’s ability to rear the animals eventually would be grossly altered by hypoxia. After the exposure, the pregnant mouse was singly-housed in a new clean cage with an intact nestlet square. The quality of the nest formed by the dam by the following morning was measured as a proxy for the dam’s ability to create a nurturing environment. Almost all mice of both conditions made typical intact nests **(Fig. 3B)** (mean/SD normoxia = 4.67 +/−0.58, hypoxia = 4.86 +/− 0.38) [43].

**Fig. 3:**
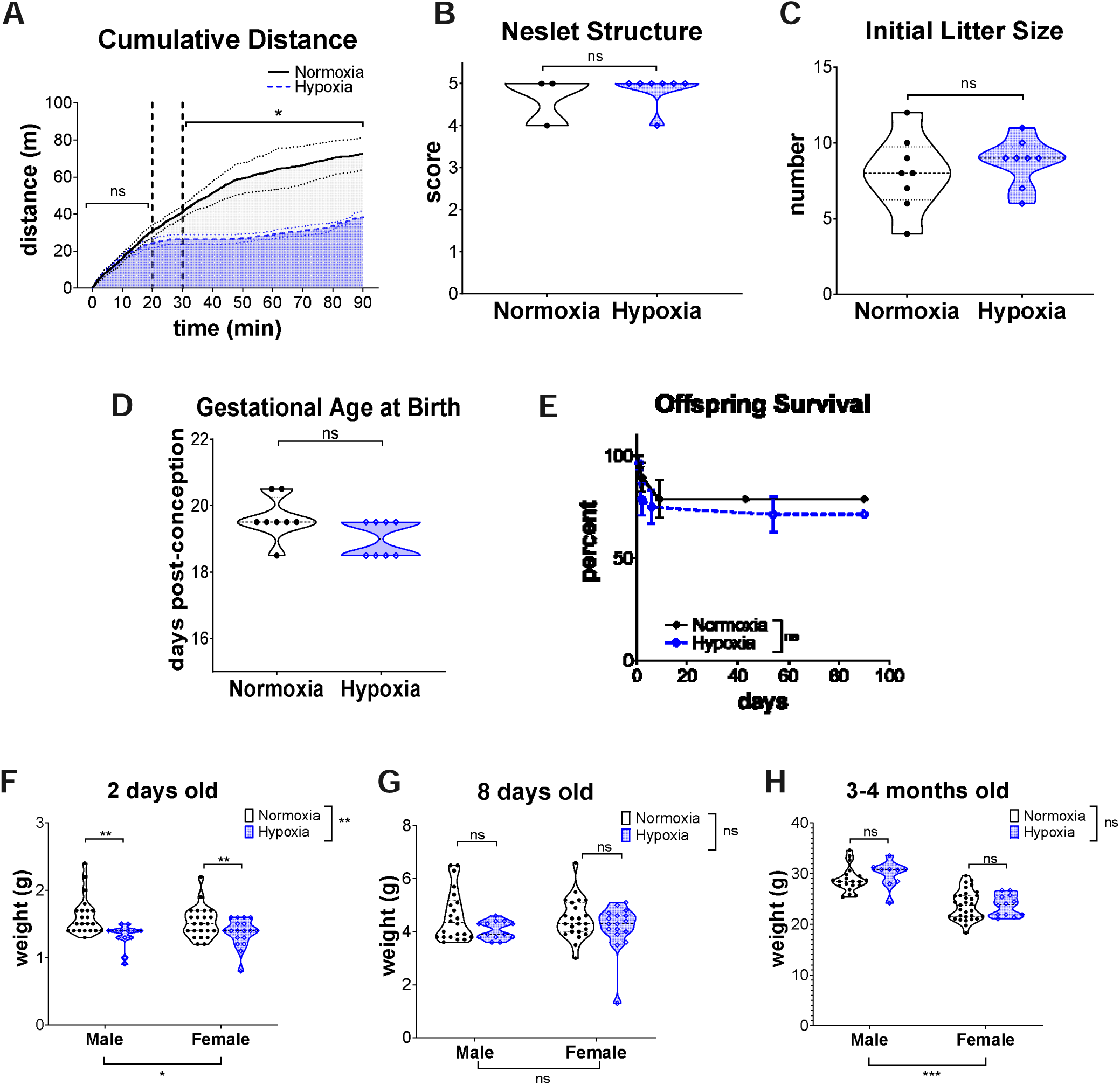
Maternal dams make typical nests after hypoxia exposure, and while there is an early decrease in weight, there are no other differences in litter health after hypoxia. (A) The cumulative distance over time of pregnant dams during the first 90 minutes the mice were in the chamber. The area under the curve is shaded. The dashed line around each line represents the SEM of each condition. The first dashed black line is where mice reach 10% inspired O2 at about 20 minutes in the chamber, and the second dashed line is when they are at the goal of 5% inspired oxygen at 30 minutes. The statistics compare normoxia and hypoxia by the area under the curve for the first 20 minutes of chamber exposure and then for 1 hour after hypoxic mice reached goal oxygen level. (B) Scoring of maternal nestlets the morning after normoxia or hypoxia. Points represent individual dams that were assessed. (C) Litter size at birth and (D) gestational age at birth for indicated individual litters were evaluated. (E) Percent survival of offspring. Error bars represent standard error at that time point. The log-rank test was used for statistics between normoxia (solid black line) and hypoxia (dashed blue line). (F-H) Weights of pups and adult offspring at the indicated age. Points correspond to the individual mice weighed. Repeated measured mixed modeling statistics are described in the methods section.

Offspring health after normoxia and hypoxia was assessed by litter size at birth, birth gestational age, survival, and weight. There was no statistical difference in litter size at birth **(Fig. 3C)** (mean/SD normoxia = 8.00 n +/− 2.45, hypoxia = 8.75 n +/− 1.58) or birth gestational age **(Fig. 3D)** (mean/SD normoxia = 19.63 days +/− 0.64, hypoxia = 19.00 days +/− 0.53). In addition, there was no difference in survival of mice through the first 90 days of life **(Fig. 3E)**. Notably, at P2 there were differences in weight due to hypoxia in males and females as well as due to sex alone **(Fig. 3F)** that were no longer present by P8 **(Fig. 3G)** (P2 weight: mean/SD normoxic male = 1.61 g +/− 0.28, hypoxic male = 1.32 g +/− 0.18, normoxic female = 1.53 g +/− 0.23, hypoxic female = 1.37 g +/− 0.21; P8 weight: mean/SD normoxic male = 4.60 g +/− 0.95, hypoxic male = 4.1 g +/− 0.34, normoxic female = 4.49 g +/− 0.76, hypoxic female = 4.17 g +/− 0.86.) There is an expected difference in weight from sex in adult animals, but hypoxia-exposure had no added effect **(Fig. 3H)** (mean/SD normoxic male = 28.87 g +/− 2.46, hypoxic male = 29.88 g +/− 2.67, normoxic female = 23.77 g +/− 2.87, hypoxic female = 23.77 g +/− 2.085). Together these results suggest that 1) maternal dams can tolerate and appropriately rear offspring after prenatal hypoxia and 2) there is no severe, life-threatening phenotype from prenatal hypoxia in exposed offspring.

### Prenatal hypoxia does not increase cell death in the fetal brain

Cell death has been noted 12-24 hours after hypoxia with ischemia in the developing brain [62, 34, 37]. Based on these findings, we isolated the fetal brain 24 hours after exposure to hypoxia and normoxia (at E18.5). We elected to use animals that had not been born yet to avoid any confounding effects of early birth on neuronal cell death [63]. A neuropathologist (A.N.V.) blinded to the exposure condition then counted apoptotic nuclei **(Fig. 4A)** in the cortex, basal ganglia, and white matter, three regions that can have significant cell death in children with acute perinatal hypoxic injury [14]. There was no increase in apoptotic nuclei in any region of the brain analyzed **(Fig. 4B–3D)** (mean/SD cortex normoxia = 1.42 cells +/− 0.52, hypoxia = 1.28 cells +/− 0.25; basal ganglia normoxia = 1.67 cells +/− 1.53, hypoxia = 0.68 cells +/− 0.58; white matter normoxia = 0.33 cells +/− 0.58, hypoxia = 0.17 cells +/− 0.28).

**Fig. 4:**
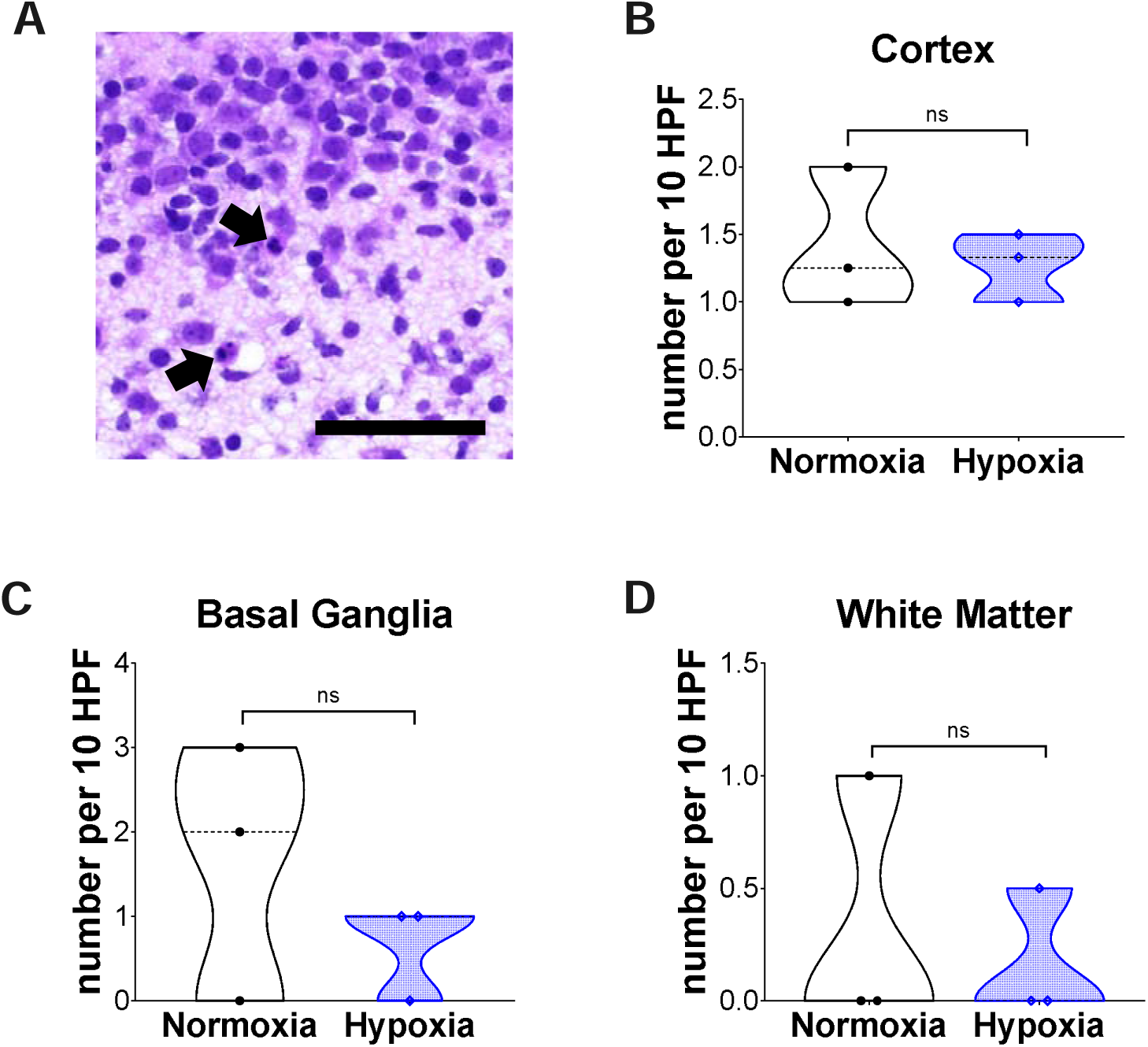
Prenatal hypoxia does not increase cell death in the fetal brain. (A) Representative image of apoptotic nuclei (arrows) within cortex, hematoxylin & eosin stain. Scale bar represents 50 μM (B-D) Number of apoptotic nuclei in the cortex, basal ganglia, and white matter reported as average seen in 10 separate high-powered fields. Statistics are described in the methods section.

### Prenatal hypoxia does not lead to long-term neuroanatomical deficits

We performed *ex vivo* MRI in adult animals to investigate potential long-term neuroanatomic deficits from late gestation transient prenatal hypoxia compared to normoxic controls. Lateral ventricle volume was measured; a single outlier had severely enlarged ventricles in the hypoxic animals **(Fig. 5A)**. However, overall, there was no significant increase in lateral ventricle size **(Fig. 5B)** (median/IQR normoxia = 1.22 mm^3^/0.33, hypoxia = 1.11 mm^3^ +/− 4.78) or differences in cortical thickness of the anterior cingulate **(Fig. 5C & D)** (mean/SD normoxia = 1.29 mm +/− 0.19, hypoxia = 1.30 mm +/− 0.09). Similarly, DTI-derived FA measurement of the corpus callosum, an index of the extent of fiber alignment and myelination, showed no difference in white matter microstructure of the genu or splenium of the corpus callosum from hypoxia **(Fig. 5C, E & F)** (mean/SD genu normoxia = 0.60 FA +/− 0.05, hypoxia = 0.59 FA +/− 0.05; splenium normoxia = 0.57 cells +/− 0.07, hypoxia = 0.59 cells +/− 0.06). We also obtained mean, axial, and radial diffusivity measurements (**Fig. 5G)**. However, processing of the post-mortem brains alters the diffusivity while FA remains unaffected [64, 65], so only the FA measurements were quantified.

**Fig. 5:**
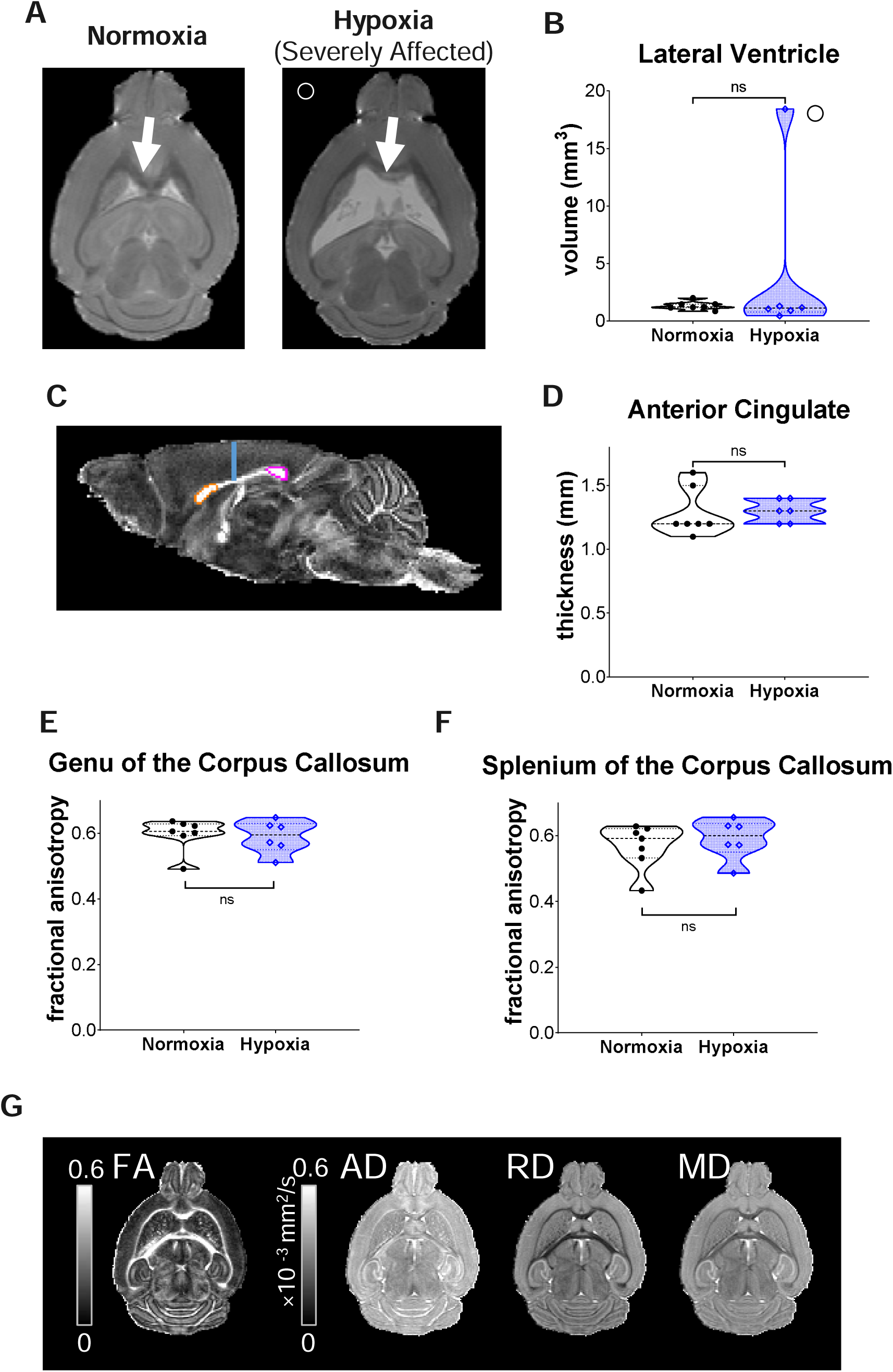
Prenatal hypoxia does not result in long-term gross neuroanatomical damage. (A) Axial images of adult female brain MRI (averaged b0 map) of indicated conditions from individual mice. (B) Quantification of ventricular size in the setting of normoxia and hypoxia. Displayed symbols in (B) represent hypoxic animals from (A). (C) Mid-sagittal image demonstrating the areas that were quantified, the anterior cingulate cortex (light blue line), the genu of the corpus callosum (orange circle), and the splenium of the corpus callosum (pink circle). (D) Quantification of cortical thickness at the anterior cingulate. (E & F) Quantification of FA in the mid-sagittal plane of the corpus callosum’s genu and splenium, respectively. Statistics as outlined in the methods. Points represent individual mice. (G) Representative image of axial section comparing FA to axial, radial, and mean diffusivity (AD, RD, and MD, respectively)

### Prenatal hypoxia leads to multiple behavior deficits in adult animals in both sexes

We tested multiple behavioral domains to determine if prenatal hypoxia leads to long-term deficits consistent with observed behavioral differences in children with prenatal hypoxic injury. Results can be divided into three categories – deficits in both males and females, sex-dichotomous deficits, and no deficits.

First, HIE is the most common cause of symptomatic neonatal seizures after delivery [66]. Weeks to years after the injury, children with moderate to severe HIE have an increased risk of developing epilepsy [67, 4]. While we did not observe any spontaneous seizures or sudden death in hypoxic animals that would be consistent with uncontrolled epilepsy, we sought to determine if hypoxic animals had a decrease in seizure threshold through flurothyl seizure threshold testing [56]. Hypoxia-exposed mice had a decreased seizure threshold (mean/SD normoxic male = 229.3 s +/− 53.5, hypoxic male = 200.0 s +/− 30.3, normoxic female = 211.8 s +/− 47.9, hypoxic female = 182.6 s +/− 27.7) **(Fig. 6A)**. These data suggest that prenatal hypoxia may predispose animals to subtle network deficits.

**Fig. 6:**
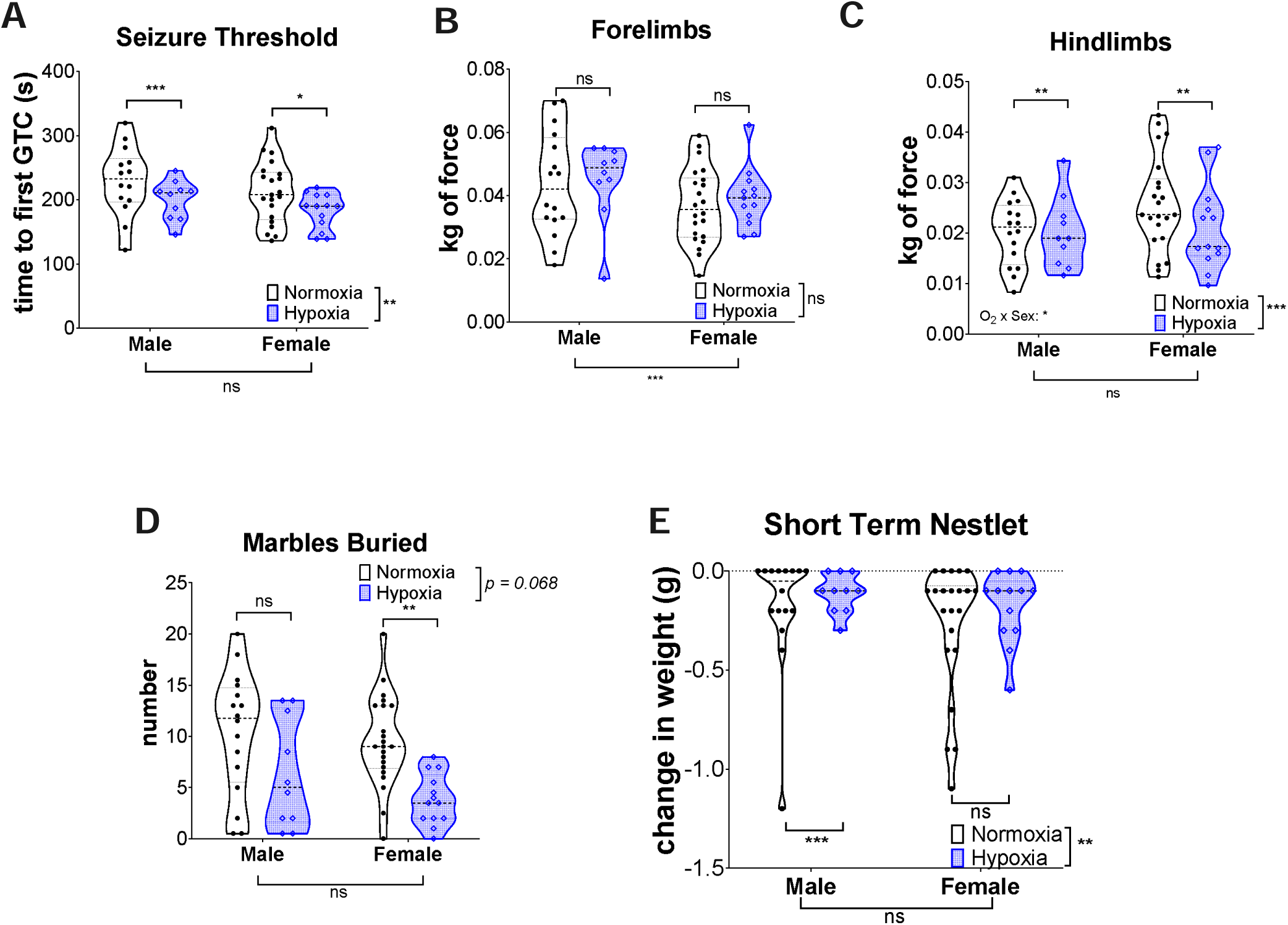
Prenatal hypoxia leads to functional deficits in adult mice. (A) Flurothyl seizure threshold study demonstrating time to first GTC. (B & C) Data from grip strength. (B) Forelimbs and (C) hindlimbs force. (D) The number of marbles buried. (E) Change in nestlet weight in short-term nestlet test. Statistics completed as outlined in the methods. Points represent individual mice.

Cerebral palsy, a non-progessive motor impairment, is also a long-term complication of HIE [68] that is sometimes correlated in animal models to abnormal grip strength [69]. We performed grip strength to determine whether hypoxic mice had long-term motor deficits [70]. Male and female animals had a baseline difference in grip strength, but there was no decrease in forelimb grip strength due to prenatal hypoxia in either sex (mean/SD normoxic male = 0.044 kg of force +/− 0.016, hypoxic male = 0.045 kg of force +/− 0.013, normoxic female = 0.037 kg of force +/− 0.012, hypoxic female = 0.039 kg of force +/− 0.009) **(Fig. 6B)**. By contrast, hypoxia-exposed mice did demonstrate significant decreases in hindlimb strength compared to normoxic control animals (mean/SD normoxic male = 0.020 kg of force +/− 0.006, hypoxic male = 0.020 kg of force +/− 0.007, normoxic female = 0.026 kg of force +/− 0.009, hypoxic female = 0.021 kg of force +/− 0.008) **(Fig. 6C)**, suggesting mild strength impairment after prenatal hypoxia.

Autism spectrum disorder has also been associated with prenatal and neonatal hypoxia [71]. Since repetitive behaviors are associated with autism, we sought to determine if there were any differences in compulsive and repetitive behaviors due to prenatal hypoxia using marble burying and short-term nestlet shredding tests [49, 51]. Increased marble burying is associated with an obsessive-compulsive related phenotype [51]. By contrast, a decrease in marble burying has been observed in genetic models of autism [72, 73]. Hypoxic animals, particularly females, had a decreased marble burying (mean/SD normoxic male = 10.34 n +/− 5.99, hypoxic male = 6.30 n +/− 5.32, normoxic female = 9.55 n +/− 4.45, hypoxic female = 3.85 n +/− 2.48) **(Fig. 6D)**. Consistent with the decrease in repetitive behaviors observed by marble burying, hypoxic mice demonstrated a significant decrease in a short-term nestlet shredding study (median/IQR normoxic male = −0.05 g /0.20, hypoxic male = −0.10 g/0.20, normoxic female = −0.10 g/0.38., hypoxic female = −0.10 g +/− 0.25) **(Fig. 6E)**. Of note, in this assay the deficit appeared more prominent in males, although there was also substantial variability in normoxic animals.

### Prenatal hypoxia leads to sex-dichotomous behavior differences in anxiety-related and social interaction-related behaviors

In addition to autism, children with HIE are also at risk for other neuropsychiatric disorders, including anxiety [71]. We used two different tests to measure anxiety-related behaviors in mice: the elevated zero maze and open field [74]. Hypoxic males demonstrated an increase in anxiety-related behaviors in both assays. The elevated zero maze test is considered a strong anxiogenic stimulus for animals, where increased anxiety is associated with decreased time in the open arms of the apparatus [74]. Hypoxic males spent less time in the open arms than normoxic males whereas hypoxia had no effect on the females (mean/SD normoxic male = 144.7 s +/− 51.8, hypoxic male = 112.0 s +/− 33.9, normoxic female = 139.0 s +/− 53.7, hypoxic female = 144.9 s +/− 40.8) **(Fig. 7A)**. We corroborated this anxiety-related phenotype in the open field test, another well-established assay for anxiety where increased anxiety is associated with decreased time in the center of the apparatus [74]. We observed that hypoxic males spent less time in the center of the field than normoxic males (median/IQR normoxic male = 151.8 s/128.8, hypoxic male = 127.6 s/105.3) **(Fig. 7B**). Females spent less time than males in the center overall, but there was no added effect from hypoxia (normoxic female = 86.3 s/69.0., hypoxic female = 84.5 s/104.7). Importantly, these differences in the open field test were not due to differences in total distance traveled (mean/SD normoxic male = 36.9 m +/− 11.5, hypoxic male = 36.6 m +/− 10.3, normoxic female = 40.8 m +/− 12.2, hypoxic female = 39.4 m +/− 13.1, *p* = 0.60 between normoxia and hypoxia).

**Fig. 7:**
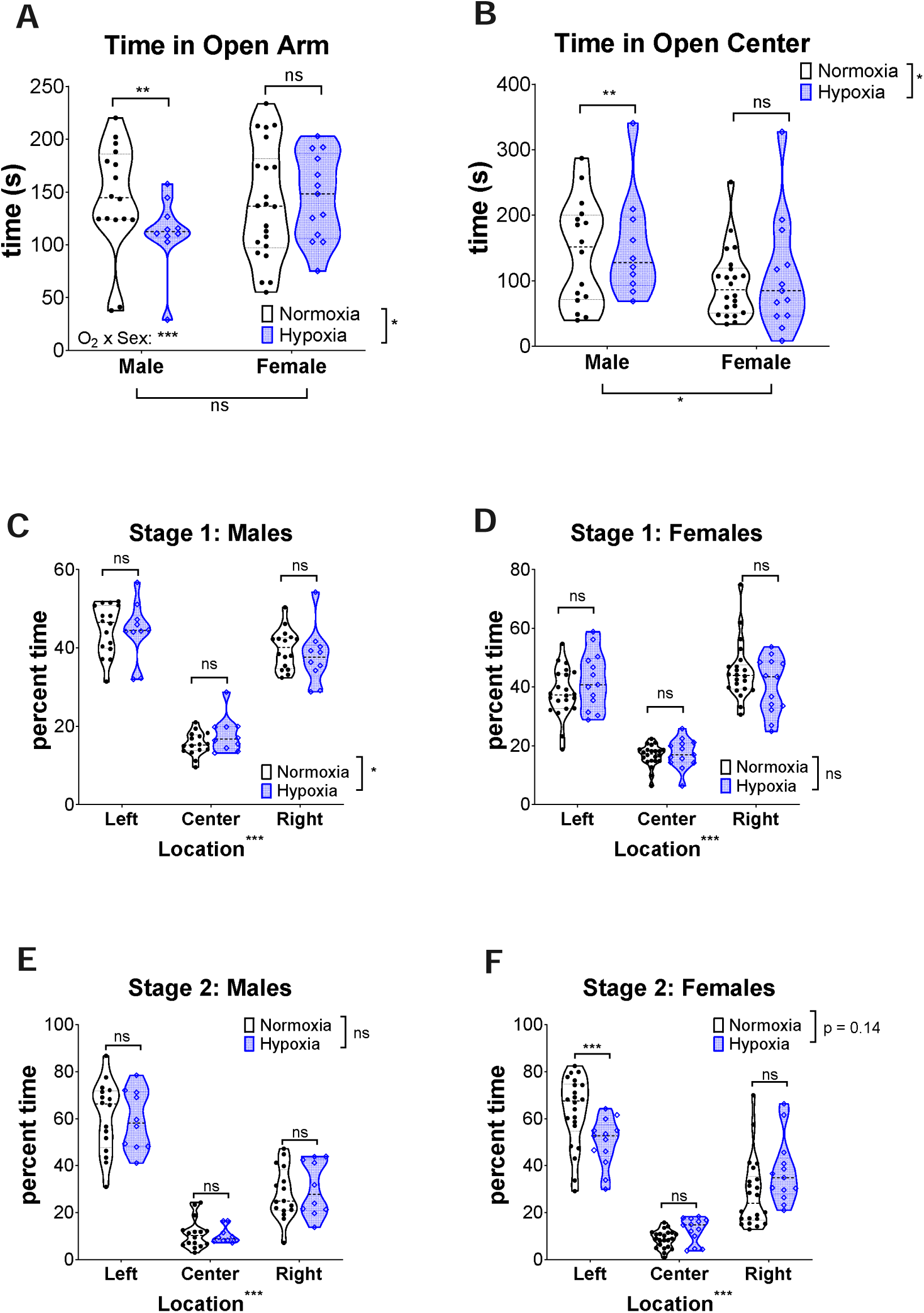
Sex-dichotomous behavior differences after prenatal hypoxia. (A) Time spent in open arms in the elevated zero maze. (B) Total time animals spent in the center of the open field. (C-F) Percent time in each mouse spent in each chamber during stages of social interaction study. (C-D) Data from habituation stage from males and females. (E-F) Data from novel mouse/object stage from males and females. Statistics as outlined in the methods. Points represent individual mice.

To further determine if prenatal hypoxia was associated with an autism-related deficit, we performed the three-chamber social interaction test. In the initial acclimation stage of this test, mice did not prefer the left or right side of the chamber **(Fig. 7C & D)** (mean/SD males normoxic: left = 44.81 m +/− 6.33, center = 15.47 m +/− 2.96, right = 39.71 m +/− 5.06; males hypoxic: left = 44.25 m +/− 7.5, center = 17.84 m +/− 4.59, right = 37.84 m +/− 7.2; females normoxic: left = 38.04 m +/− 8.30, center = 16.65 m +/− 3.69, right = 45.28 m +/− 9.69; females hypoxic: left = 42.24 m +/− 9.66, center = 17.28 m +/− 5.03, right = 40.44 m +/− 9.55). In the second stage, a novel mouse was placed in a cup in the left side of the chamber and a novel object was placed in the right. As expected, normoxic males and females preferred the novel mouse to the novel object **(Fig. 7E & F)**. Hypoxic males demonstrated a similar increase in preference for the novel mouse to normoxic controls **(Fig. 7E)** (mean/SD males normoxic: left = 60.96 m +/− 15.16, center = 11.01 m +/− 6.31, right = 28.02 m +/− 11.00; males hypoxic: left = 59.35 m +/− 12.71, center = 10.30 m +/− 3.48, right = 30.35 m +/− 11.62). Hypoxic females, however, had a weaker preference for the novel mouse than the normoxic controls **(Fig. 7F)**, suggesting a hypoxia-related decrease in social interaction only in female animals (mean/SD females normoxic: left = 63.13 m +/− 14.92, center = 8.74 m +/− 3.80, right = 28.12 m +/− 14.84; females hypoxic: left = 50.06 m +/− 10.27, center = 12.61 m +/− 5.31, right = 37.33 m +/− 13.70).

### Prenatal hypoxia does not lead to deficits in learning or memory

To determine whether prenatal hypoxia led to deficits in learning or memory, we tested motor learning and visual-spatial learning and memory. Rotarod testing for motor learning and coordination [53] showed no difference between hypoxia and normoxia **(Fig. 8A**).

**Fig. 8:**
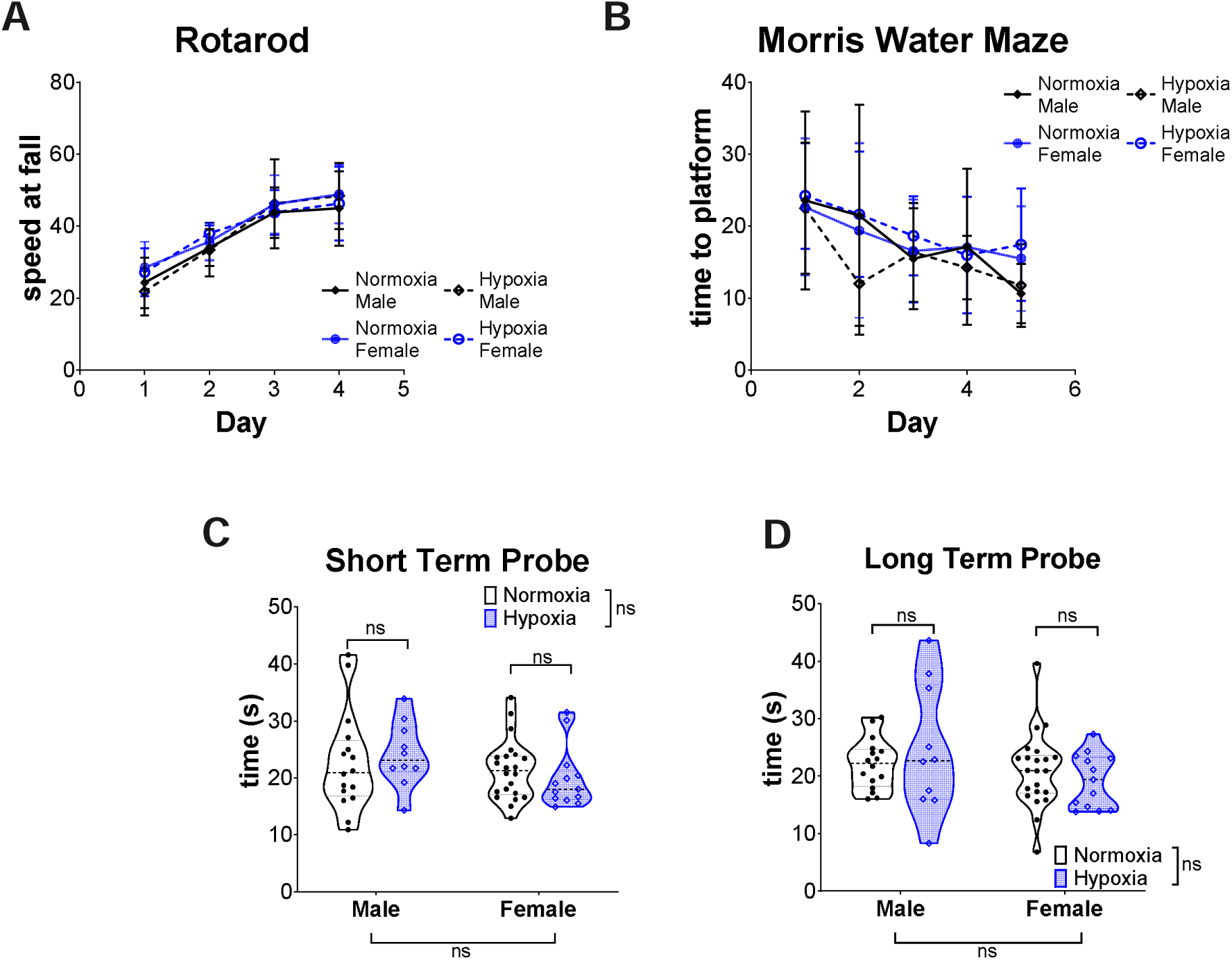
Prenatal hypoxia does not lead to deficits in learning or memory. (A) The speed at fall on Rotarod. (B-C) Data from the Morris Water Maze. (B) Latency time to platform in place trials. (C) One-hour probe trial. (D) Twenty-four-hour probe trial. Statistics were performed as detailed in methods. Each point represents a single animal.

The Morris water maze was used to test for deficits in learning and memory [50]. There was no difference between normoxic and hypoxic mice during the place trials in time to the platform **(Fig. 8B)**. Probe testing in the Morris water maze was used to identify any differences in memory. Prenatal hypoxia did not predispose mice of either sex to deficits in short or long-term memory **(Fig. 8C & D)** (mean/SD short-term probe normoxic male = 22.73 s +/− 8.68, hypoxic male = 24.12 s +/− 5.69, normoxic female = 21.40 s +/− 5.27, hypoxic female = 19.86 s +/− 5.29; long-term probe normoxic male = 22.11 s +/− 4.40, hypoxic male = 24.48 s +/− 11.17, normoxic female = 21.00 s +/− 6.57, hypoxic female = 19.24 s +/− 4.69). Together, these data suggest that this prenatal hypoxia does not predispose animals to significant learning or memory deficits.

## Discussion/Conclusion

Here we describe that in a simple prenatal hypoxia mouse model, the animals have a molecular response to hypoxia in the fetal brain and some evidence of long-term functional deficits despite not having any early cell death or long-term gross neuroanatomic injury. Some of these deficits are analogous to NDDs seen in children years after HIE injury, including deficits in anxiety [75], motor function [76], and susceptibility to seizures [4, 77]. In addition, the functional deficits in the absence of long-term structural deficits seen in this model are also consistent long-term NDDs seen in children with mild HIE who also lacked significant evidence of structural brain injury [4, 78]. However, this model did not produce significant deficits in motor learning or visual-spatial learning that more directly correlate to cognitive deficits seen in these children [7]. Together, these data suggest that this model is amenable for understanding the contribution of prenatal hypoxia to long-term deficits in functional outcomes that can occur in the absence of significant structural damage.

This model provides a clinically-relevant paradigm for investigating mild and preterm HIE through an *in utero* injury. One of the strengths of this model is that, like most incidences of HIE in neonates, it is a prenatal injury and, therefore, can be used to understand the physiology of *in utero* insult. It is crucial to understand the specific effects of pathologic levels of hypoxia in the *in utero* environment because the fetal environment is intrinsically hypoxic at baseline [20, 35], which has implications in energy metabolism and how the fetal brain may compensate for pathologic hypoxic exposure. For instance, the fetal brain is already more likely to use anaerobic respiration than the postnatal brain for energy [79], indicating a possible lower tolerance for a hypoxic insult if the fetal brain is driven further away from aerobic respiration. Supporting the hypothesis that *in utero* brain metabolism status may be important for understanding HIE severity, antenatal risk factors that lead to placental insufficiency, like excessive gestational weight gain, portend to worse outcomes from HIE [28]. This model is poised to study these acute-on-chronic interactions in a manner that is impossible to investigate in postnatal models.

Previous investigations informed the parameters for hypoxic exposure we used in this study [20, 27]. Photoacoustic ultrasound experiments demonstrate that 5% inspired oxygen reduces blood flow to the uterus and placenta within minutes [27]. This study also noted that the placenta had a significant reserve compared to the maternal skin to withstand substantial transient changes in oxygen levels [27]. Therefore, we opted for prolonged hypoxia exposure in this study to overcome this reserve. Uterine and placental blood flow studies would have required a pregnant mouse to be anesthetized for the 8 hours of the exposure, which is not feasible. We reasoned that since the rest of the study focused on outcomes of animals exposed to fetal hypoxia, we wanted to directly test the effects of hypoxia on the organ of interest – the fetal brain. To determine as directly as possible if the fetal brain responded to hypoxia, we relied on the well-known hypoxia-response pathway of Hif1α and *Vegfa*. This study reproduced findings that Hif1α protein is stabilized in the fetal cortex of normoxic animals [20]. We further demonstrated increased expression of *Vegfa*, a downstream Hif1α target, within the first 4 hours of hypoxia. Hif1α is involved in metabolic regulation [80] and is required for neuronal and glial development [21, 81, 22], indicating the importance of regulating the oxygen response for cell survival and differentiation in the developing brain.

Interestingly, Hif1α levels peaked at 4 hours of hypoxia and then decreased at 8 hours. Provocatively, this decrease suggests that there may be an initial capacity to compensate for hypoxia that is lost over time. The amount of time *in utero* hypoxia needed for children to experience brain injury is elusive because the diagnosis of HIE relies on recognizing the hypoxic event by the mother or healthcare providers [14, 82]. It is possible that in some instances, there is a more prolonged, more indolent hypoxic insult before there is a clinical concern for HIE. Therefore, it is possible our observation that Hif1α levels change at different durations of hypoxia may someday be clinically relevant for understanding the kinetics of an injury. One potential therapeutic application of this finding is that suppose future studies using this model of prenatal hypoxia discover hypoxic preconditioning can protect exposed offspring from deficits as is seen in postnatal models [25, 26]. In that case, the dynamics of *in utero* Hif1α stabilization after hypoxia may be necessary for understanding the mechanisms underlying this protective effect.

This model may also prove helpful in studying the effects of sex on outcomes from prenatal hypoxia. Clinical studies of long-term outcomes of HIE are underpowered to determine sex-related differences in anxiety or cognition in school-aged children [83, 77]. However, we hypothesize that children with HIE have sex-dichotomous differences in outcomes, as is seen in children with brain injury from prematurity or intraventricular hemorrhage [40]. Postnatal models of hypoxia and hypoxic with ischemia have sex differences in several behaviors [39–42], supporting this hypothesis. At present, though, it is unclear if the sex-specific differences in anxiety-like behaviors or social interactions we found in this model are clinically relevant. In particular, in this study, females were more likely to have subtle deficits in social interaction, but, in children, boys are more likely to be affected by autism [84]. Adding additional complexity to this question, sex differences in behavior may fluctuate throughout social maturation [85], and there are sex-dependent regional differences in brain development [86]. The MRI studies presented here are underpowered to determine if hypoxia has subtle sex-dichotomous effects on ventricle size, cortical thickness, or volumes of specific regions.

One of the caveats of this model requires understanding the differences between human and mouse brain development **(Fig. 1A)**. The rodent brain at postnatal day 0 is similar to a 23-26 week gestation human brain and is considered human “term” equivalent at postnatal days 7 to 10 [16]. Therefore, this model is most suited for studying the neuroanatomic changes expected from preterm HIE. White matter microstructure is affected in premature infants [87, 88], but we did not see differences in the selected ROIs of the corpus callosum (e.g., genu or splenium). However, the lack of abnormality in the corpus callosum does not rule out more subtle effects in white matter development. *Ex vivo* MRI studies with more animals may help identify subtle differences in white matter microstructure due to prenatal hypoxia. Similarly, cortical thickness or increased *ex vacuo* ventriculomegaly is unlikely to be sufficiently sensitive to capture more subtle changes in neuronal structure. However, further histological assessments may reveal cell type-specific effects of hypoxia.

In addition, some curious differences exist between the acquisition of neurodevelopment milestones and functional networks between humans and rodents [30]. For example, mice do not exhibit respiratory distress or feeding dyscoordination at birth which is frequently present in children born at less than 34 weeks gestation [89]. Furthermore, mice acquire the ability to crawl within a few days of life and are generally capable of immature walking by postnatal day 10, which is typically a developmental milestone achieved by a 12-month-old infant [30]. Thus, future studies will use this model of prenatal hypoxia to investigate the effect of hypoxia on these clinically relevant neurodevelopmental milestones that are still developing in the term human brain.

A limitation of this study is that we only performed the behavior and imaging studies at a single time point in adult animals. Animals demonstrate significant changes in behavior and brain structures as they age and undergo sexual maturation [90, 91, 85, 92]. By selecting this time point, we did not confound the initial characterization of this model with the effects of concurrent sexual maturation. However, we may have missed changes from hypoxia that were only present in juvenile animals in both behavior and structure. We hope future studies using this model can focus on the potential relationship between particular brain regions and sex-dichotomous behavioral differences throughout maturation.

Another limitation of this model is that it does not have a proper “ischemic” component since we did not actively clamp blood flow to the developing brain. We likely did not observe more severe behavioral deficits and increased cell death due to the lack of ischemia. In the Vannucci model, animals demonstrate visual-spatial learning deficits [15] that are absent in this model. We postulate these deficits in the Vannucci model are likely related to the extent of hippocampal cell death from the carotid ligation, not solely to the differences in gestational age. Supporting this theory, a mouse model of transient uterine clipping at E16.5 demonstrated increased cell death 12 hours after the occlusion and significant cognitive deficits [34]. Studies in prenatal sheep models also support that hypoxia and ischemia, but not hypoxia alone, are required for extensive cell death [37, 38]. These differences in cell survival between hypoxia only and hypoxia with ischemia are replicated in cell culture. Neurons survive in hypoxia alone but are much more susceptible to cell death in the setting of hypoxia with glucose depletion [93]. In a human spheroid model of hypoxic injury from prematurity, there is no significant increase in cell death after 48 hours of 1% inspired oxygen [94]. Therefore, our finding of no cell death is consistent with the literature. However, like transient prenatal hypoxia in sheep, we also demonstrate long-term functional deficits, most notably in seizure threshold differences. Thus, this model can still be leveraged to understand the molecular mechanisms driven primarily by hypoxia, leading to functional deficits.

In conclusion, we present a simplified model of prenatal hypoxic insult that phenocopies many functional deficits consistent with findings in children with mild HIE. We expect these results will provide a platform to study the effects of prenatal risk factors and the sex-dichotomous impact of hypoxia. In addition, this prenatal model may offer an attractive platform for studying the genetic factors that exacerbate the mild phenotype (e.g., determine which factors protect from cell death in the setting of hypoxia only). In the end, with comparative studies amongst prenatal and postnatal injury models, we can better understand the full spectrum of relevant NDDs from hypoxia to the developing brain and determine opportunities for novel interventions.

## Acknowledgments

We thank the Small Animal Imaging Facility at the Research Institute of the Children’s Hospital of Philadelphia (CHOP), the CHOP Pathology Core, the Neurobehavior Testing Core, and EEG./Epilepsy core at UPenn and IDDRC at CHOP/Penn [Grant: U54 HD086984] for assistance. We thank Dr. Rui Xiao in the Department of Biostatistics at UPenn for her advice on statistical analysis. We thank Brenna Daugherty for initial studies monitoring pregnant mouse locomotion. We appreciate Daniel Licht and the Wolfson family for providing the Biospherix oxygen chambers. Finally, we are thankful to Dr. Michal Elovitz and members of the Marsh and Anderson labs for numerous insightful discussions on the data in this manuscript.

## Statement of Ethics

Studies involving animals were approved by the Institutional Animal Care and Use Committee (IACUC) at the Children’s Hospital of Philadelphia (Approved Number: IAC 22-000547).

## Author Contributions

Ana G. Cristancho conceived of the project designed and performed, analyzed, and interpreted components of all the experiments. Elyse C. Gadra completed immunoblots and behavior experiments. Ima M. Samba conducted behavior experiments and assessed maternal dam health. Chenying Zhao and Sergey Magnitsky performed the MRI experiments. Minhui Ouyang and Hao Huang analyzed the MRI experiments. Angela N. Viaene completed the histology for apoptosis. Stewart A. Anderson and Eric D. Marsh participated in conceiving the project and interpretation of the data. Ana G. Cristancho wrote the initial manuscript and finalized all versions. Ana G. Cristancho, Elyse C. Gadra, Ima M. Samba, Chenying Zhao, Minhui Ouyang, Sergey Magnitsky, Hao Huang, Angela N. Viaene, Stewart A. Anderson, and Eric D. Marsh participated in revisions of the manuscript.

## Data Availability Statement

All data generated or analyzed during this study are included in this article. Further inquiries can be directed to the corresponding author.

